# Explorative meta-analysis of 417 extant archaeal genomes to predict their contribution to the total microbiome functionality

**DOI:** 10.1101/2020.08.04.236075

**Authors:** Robert Starke, Maysa Lima Parente Fernandes, Daniel Kumazawa Morais, Iñaki Odriozola, Nico Jehmlich, Petr Baldrian

## Abstract

Unveiling the relationship between taxonomy and function in microbiomes is crucial to determine their contribution to ecosystem functioning. However, while the relationship between taxonomic and functional diversity in bacteria and fungi was reported, this is not the case for archaea. Here, we used a meta-analysis of completely annotated extant genomes of 417 taxonomically unique archaeal species to describe intergenome and intragenome redundancy of functions and to predict the extent of microbiome functionality on Earth contained within archaeal genomes using accumulation curves of all known functions from the level 3 of KEGG Orthology. We found that intergenome redundancy as functions present in multiple genomes was inversely related to intragenome redundancy as multiple copies of a gene in one genome, implying the trade of between additional copies of functionally important genes or a higher number of different genes. A logarithmic model described the relationship between functional diversity and species richness better than both the unsaturated and the saturated model, which suggests a limited total number of archaeal functions in contrast to the potential of bacteria and fungi. Using a global archaeal species richness estimate of 13,159, the logarithmic model predicts a total of 4,164.1 ±2.9 KEGG level 3 functions while the non-parametric bootstrap estimate yields a lower bound of 2,994 ±57 KEGG level 3 functions. Our approach not only highlights similarities in functional redundancy but also the difference in functional potential of archaea compared to other domains of life.

## Introduction

Ecosystem functioning is mediated by biochemical transformations performed by a community of microbes from all domains of life (Woese et al., 1990). Even though ecological studies tend to focus on bacteria and fungi, archaea as major part of global ecosystems (Delong, 1998) are ubiquitous in both terrestrial and aquatic environments (DeLong and Pace, 2001; Timonen and Bomberg, 2009). Particularly, archaea make up between 20% and 30% of the total prokaryotes in marine environments (Stoica and Herndl, 2007) and between 0% and 10% in soil environments (DeLong and Pace, 2001; Bates et al., 2011). Increases in archaeal abundance were found in extreme niches such as acidity and low temperature (Korzhenkov et al., 2019). Functionally, archaea play key roles in global carbon (e.g., methanogenesis or CO_2_ fixation) and nitrogen (e.g., N_2_ fixation or oxidation of ammonia) cycles (Leininger et al., 2006) but they also have complex relationships with both bacteria and fungi (Bengtson et al., 2012). In every community, multiple organisms from different taxonomic groups can play similar if not identical roles in ecosystem functionality, the so-called functional redundancy (Hubbell, 2005). In fact, interspecies redundancy of certain functions was shown to be very high with several hundreds to thousands of different taxa expressing the same function within one habitat (Žifčáková et al., 2017). These functions can be statistically inferred based upon homology to experimentally characterized genes and proteins in specific organisms to find orthologs in other organisms present in a given microbiome (Starke et al., 2019, 2020b, 2020a). This so-called ortholog annotation is performed in KEGG Orthology (Kanehisa et al., 2016a, 2016b) that covers a wide range of functional classes (level 1 of KEGG) comprising cellular processes, environmental information processing, genetic information processing, human diseases, metabolism, organismal system, *brite* hierarchies and functions not included in the annotation of the two databases *pathway* or *brite* (more information about the databases can be obtained under https://www.genome.jp/kegg/kegg3.html). However, the bottleneck of describing microbiome functions is the low number of fully sequenced and annotated genomes as they are mostly limited to those that have undergone isolation and extensive characterization (Starke et al., 2019, 2020b, 2020a) with respect to the expected total diversity. Hence, the lower the share of known species or the higher the predicted total diversity, the weaker is the prediction itself. Problematically, the vast majority of organisms were not yet studied (Pham and Kim, 2012; Martiny, 2019) and the annotation is based on the similarity to the genomes of the very few studied model organisms (Starke et al., 2019, 2020b, 2020a). As a consequence, microbiome functionality can be inferred based on its taxonomic composition and its relation to functional parameters (Starke et al., 2018) as indicated by the frequent use of nuclear ribosomal 16S metabarcoding to describe the prokaryotic community. Although the description of microbial communities is important to assess the drivers of the occurrence of individual taxa and the composition of their communities (Starke et al., 2020b), the mere community composition in itself does not provide detailed answers, i.e. about its functional diversity (VětrovskÝ et al., 2019). The functional diversity for both bacteria (Starke et al., 2020a) and fungi (Starke et al., 2019, 2020b, 2020a) were recently predicted to comprise millions of different functions using meta-analyses of proteins (Starke et al., 2020a) and genomes (Starke et al., 2019, 2020b); most of which are unknown today. However, our understanding of functional redundancy in archaea and their contribution to the total microbiome functionality is still scarce.

Here, we used both parametric and non-parametric estimators of functional richness with the aim to predict the total archaeal functionality on Earth and to unveil the relationship between taxonomy and function in the archaeal domain. For this, we extracted all completely annotated genomes of taxonomically unique archaeal species (n=417) from the Integrated Microbial Genomes and microbiomes (IMG) of the Joint Genome Institute (JGI) (https://img.jgi.doe.gov/) on July 17^th^ 2020 with taxonomic annotation on species level and functional annotation on level 3 of KEGG. We analyzed the relationship of gene counts and number of KEGG functions within the archaeal domain and used a parametric estimation based on an accumulation curve (AC) (Gotelli and Colwell, 2001) characterized by increasing number of KEGG level 3 functions with increasing species using 1,000 random permutations and its subsequent fit to a saturated, an unsaturated and a logarithmic model. Chao-1 for every 50 randomly picked species of all 417 in the database each with 20 replicates represented the non-parametric estimator. The precision of both the parametric and the non-parametric approach generally depend on the proximity to the asymptote of the model; with greater extrapolation to the total count resulting in greater error (Thompson et al., 2003). We therefore hypothesized more precise estimates of the contribution of archaea to the total microbiome functionality than previously calculated for both bacteria (Starke et al., 2020a) and fungi (Starke et al., 2019, 2020b, 2020a) due to the higher coverage of predicted taxonomic diversity of archaea.

## Results

### Gene counts and number of KEGG level 3 functions

The gene count per genome in archaeal phyla was significantly higher (HSD-test) in *Euryarchaeota* as compared to *Crenarchaeota* and *Thaumarchaeota* (**Figure 1a**). On the level of habitats, archaea isolated from fresh water, sediments or soils had on average significantly more genes than archaea enriched from the deep sea or hot springs. A comparable number of archaeal genomes were sampled from each habitat, ranging from 8 in sludge to 37 in hot springs. On the level of temperature preferences, mesophilic archaea comprised significantly (HSD-test) more genes than thermophilic and hyperthermophilic archaea. Similar significant differences were found in the number of KEGG level 3 functions on all three prior investigated levels (**Figure 1b**).

**Figure 1:**
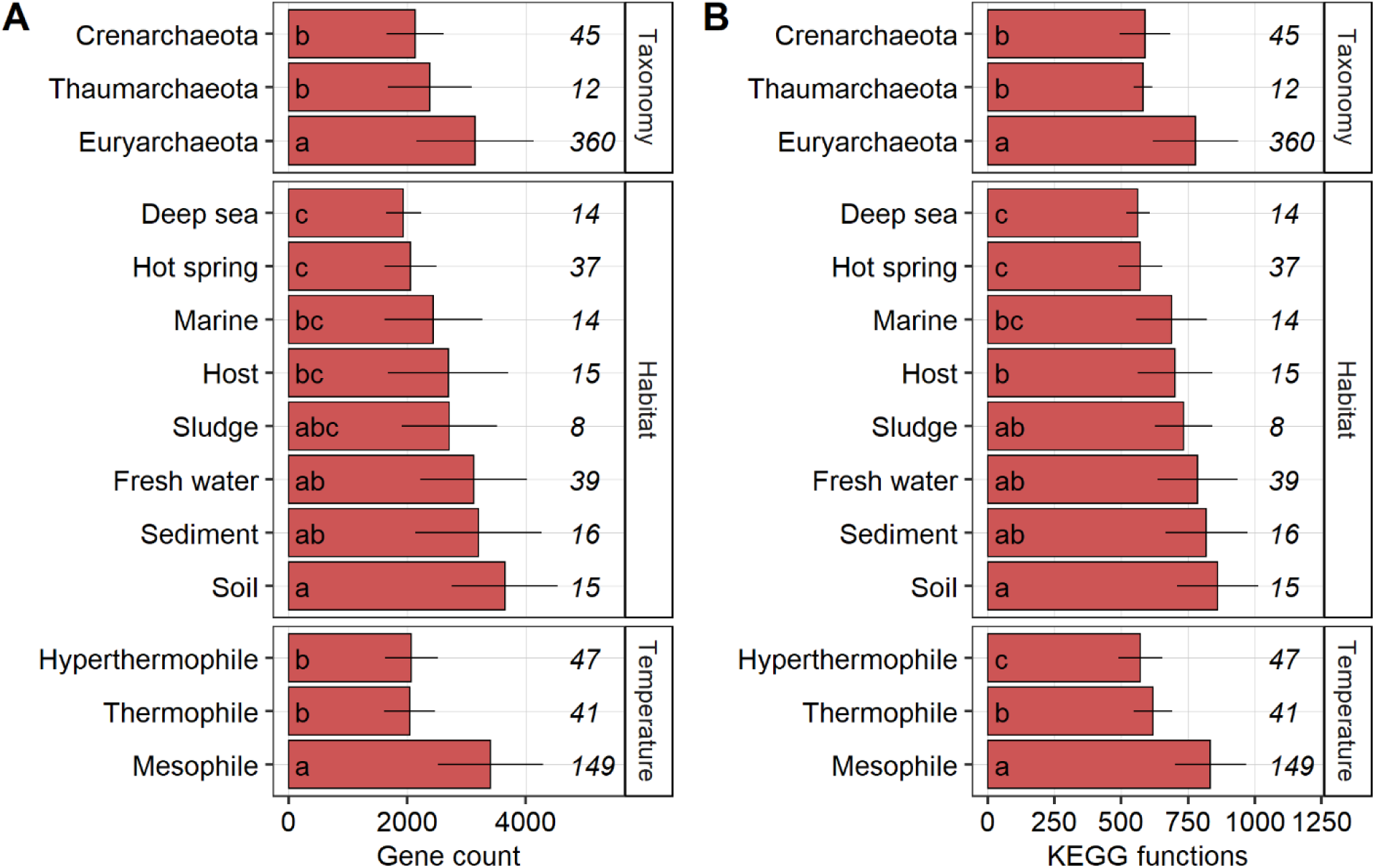
Gene counts (a) and the number of different KEGG functions (b) per genome across archaeal phyla, habitats and temperature ranges shown as average with standard deviation. The number of archaeal genomes is given in italic. Groups followed by a different letter are significantly different according to the HSD-test (*P* < 0.05).

### Inter- and intragenome functional redundancy

Intergenome functional redundancy describes the performance of one metabolic function by multiple taxonomically distinct organisms while intragenome functional redundancy accounts for the number of replicated functions within one genome (Starke et al., 2019, 2020b). Across all 417 archaeal genomes, the median of intergenome functional redundancy was found to be 0.06 (**Figure 2a**). The majority of functions were found with low redundancy as 1,650 KEGG functions were found in less than 10% of the species. In comparison, only 172 KEGG functions were found in more than 90% of the archaeal genomes. Together, 65.3% of all functions showed either high or low redundancy while the rest appeared intermediate with an intergenome functional redundancy between 0.1 and 0.9 with a particularly high abundance at around 0.24. The median of intragenome functional redundancy across all 417 archaeal genomes was found to be 1.02 gene copies per KEGG function with a maximum of 72 gene copies (**Figure 2b**). Among archaeal phyla, *Thaumarchaeota* showed a significantly higher (HSD-test) intergenome functional redundancy compared to *Crenarchaeota* and *Euryarchaeota* (**Figure 3a**). Within habitats, the intergenome redundancy in the deep sea, hot springs, sediments and sludges were significantly higher (HSD-test) than in fresh water, host, marine and soil habitats. On the level of temperature preferences, intergenome redundancy was highest in hyperthermophilic archaea, followed by thermophilic and mesophilic ones. The inverse pattern was found for intragenome functional redundancy for all three investigated levels (**Figure 3b**). Significantly higher intergenome redundancy was accompanied by significantly lower intragenome redundancy and vice versa regardless the taxonomy, habitat and temperature preference of archaea.

**Figure 2:**
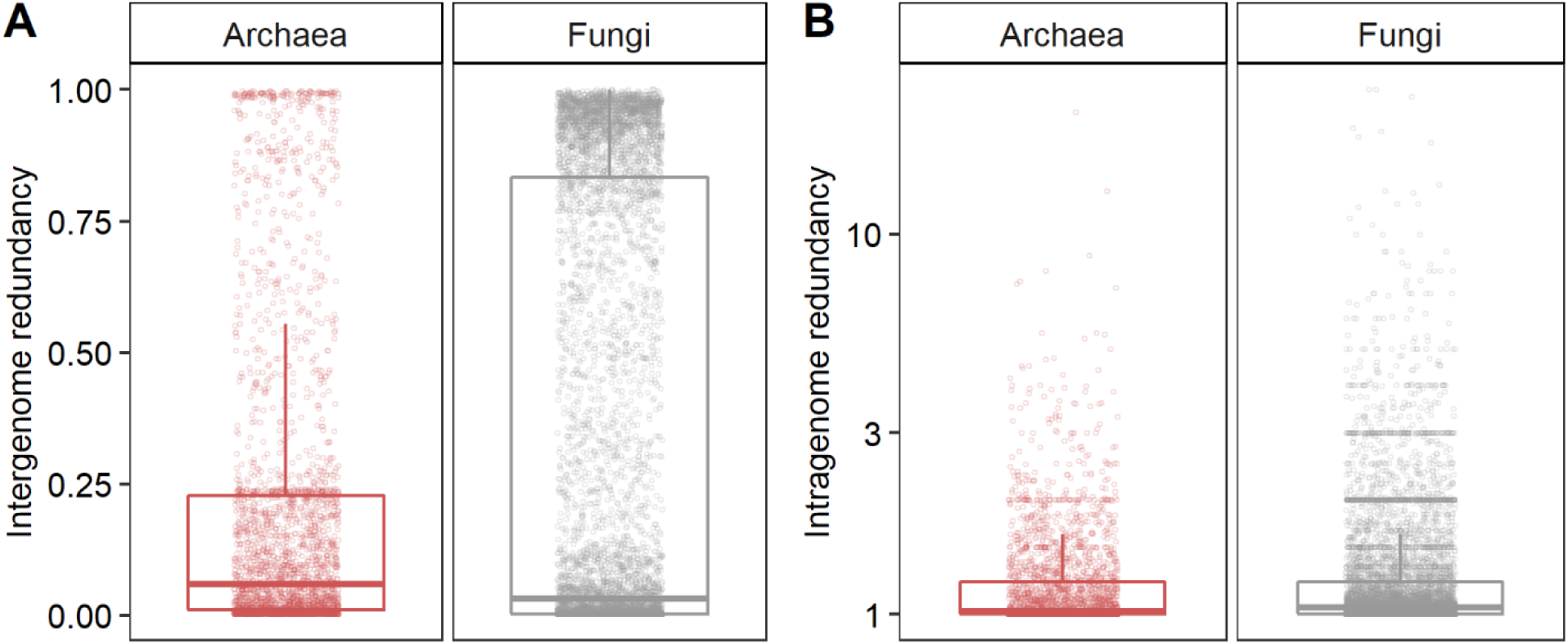
The distribution of intergenome functional redundancy as the total share of functions within archaea relative to the total number of archaeal species in the database (a) and intragenome functional redundancy as the number of replicated KEGG functions within one archaeal species in the database (b) compared to the previously published distributions in fungi (Starke et al., 2020b).

**Figure 3:**
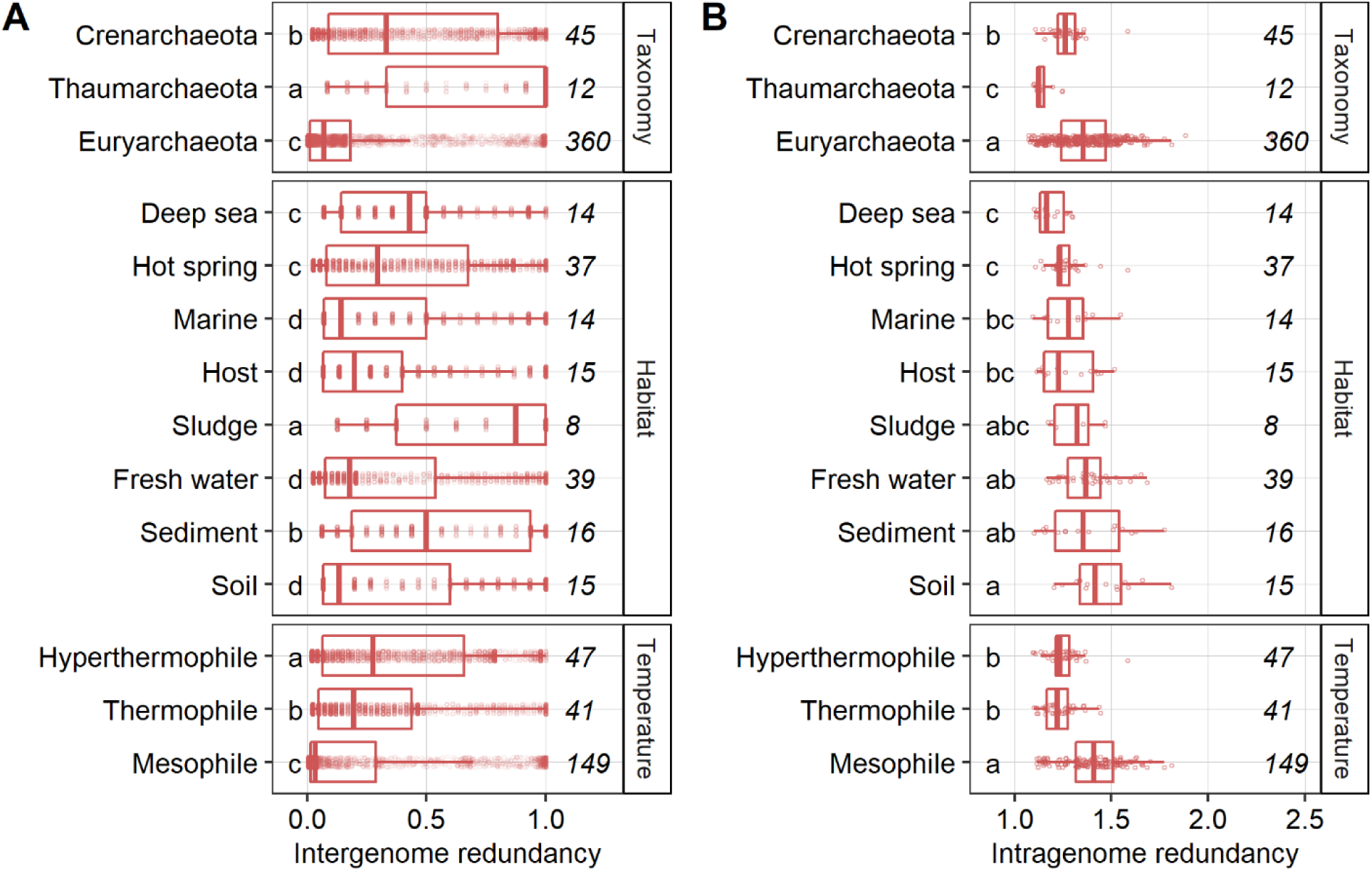
Intergenome (a) and intragenome functional redundancy (b) in archaeal phyla, habitats and temperature ranges. The number of archaeal genomes is given in italic. Groups followed by a different letter are significantly different according to the HSD-test (*P* < 0.05).

### Parametric and non-parametric estimation of the archaeal contribution to the total microbiome functionality

The logarithmic model described the dependence of functional categories on species richness significantly better than both the saturated and the unsaturated model, estimated by lower Akaike Information Criterion (AIC) (**Figure 4a**), to imply a plateau of functional richness with higher species richness. Considering the estimate of 13,159 archaeal species on Earth (Yarza et al., 2014) and assuming that the relationship between species richness and functional richness will be logarithmic with more species, we propagated the logarithmic model with the result of a total archaeal functionality of 4,164.2 ±2.9 KEGG functions (with 4,158.6 and 4,169.9 as 95% confidence intervals). Similarly, the non-parametric estimator of functional richness, assuming the existence of a maximum functional richness, indeed plateaued for the 417 archaeal genomes (**Figure 4b**). Estimations obtained with more than 200 archaeal genomes generated broadly overlapping confidence intervals indicative of the reliability of the estimation of the asymptotic functional richness. The three non-parametric estimators yielded comparable estimations of asymptotic functional richness: 3,128 ±42 KEGG functions using the Chao-1 index, 3,169 ±78 using the first order jackknife and 2,994 ±57 using the bootstrap method (**Figure 4b**).

**Figure 4:**
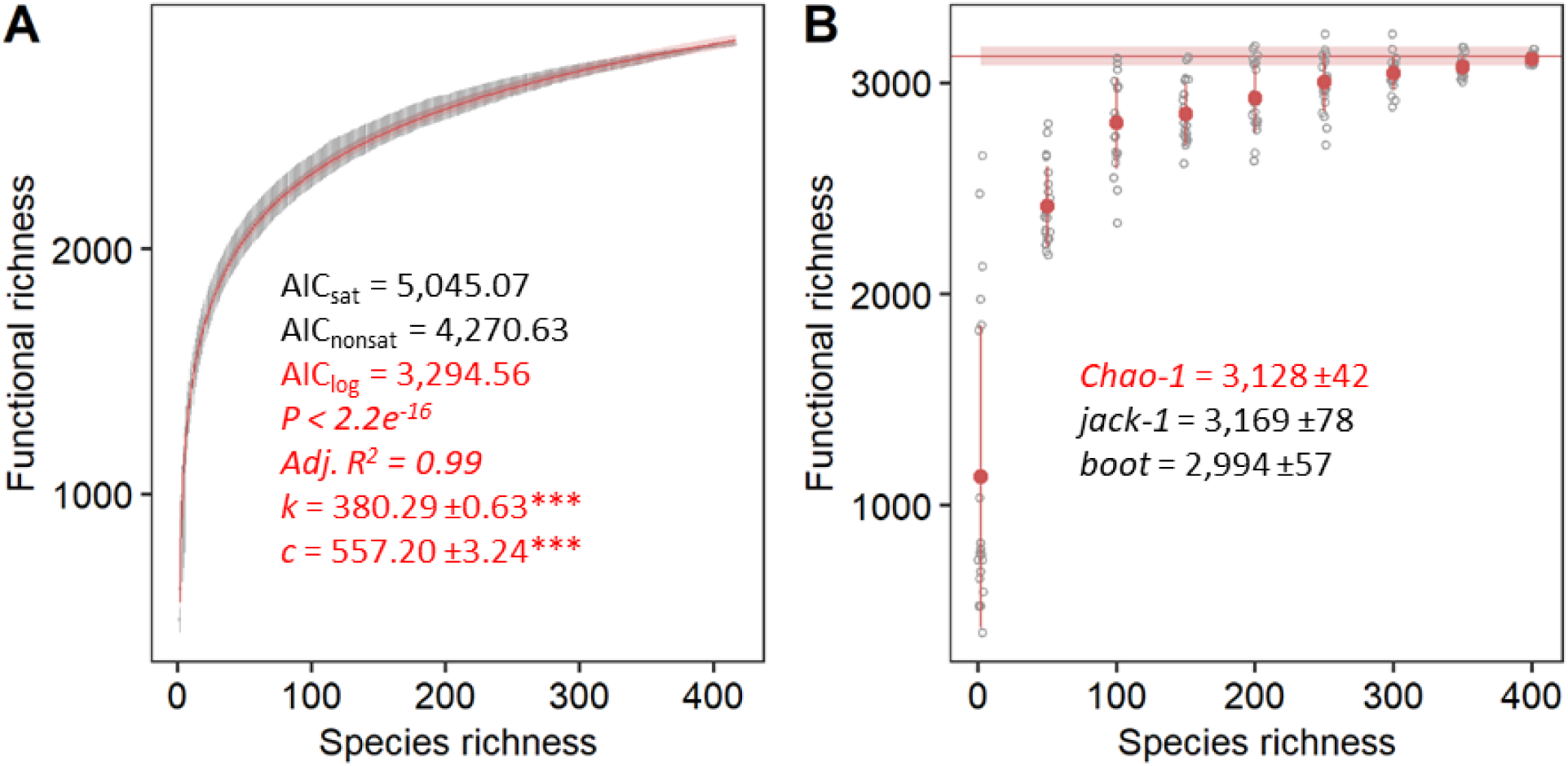
Parametric (a) and non-parametric (b) estimation of the total functional richness. The logarithmic model of the accumulation curves as grey points with error bars as 95% confidence intervals for the total known archaeal microbiome functions derived from the KEGG database by 1,000 random permutations for every one species richness with 95% confidence intervals. Significance of the parameter estimates are indicated by asterisks (*** equals *P* < 0.001). The Chao-1 index as lower bound and non-parametric estimate was calculated using 20 replicates shown in grey for every 50 randomly picked archaeal genomes in the database starting with two species. The Chao-1 index of all 417 archaeal genomes with standard errors is shown as red line.

## Discussion

### Genome content

Generally, the genome size of an organism depends on its developmental and ecological needs (Petrov, 2001). A large genome directly increases both nuclear and cellular volumes (Cavalier-Smith, 1978), which helps to buffer fluctuations in concentrations of regulatory proteins or to protect coding DNA from spontaneous mutation (Vinogradov, 1998). Hence, variation in genome size is due to adaptive needs or due to natural selection in different organisms (Petrov, 2001); the so-called adaptive theory of genome evolution. A higher evolutionary rate (Wang et al., 2010) could be directly related to the small genomes of archaea. The 417 archaeal species generally comprised of smaller genomes compared to bacteria (VětrovskÝ and Baldrian, 2013) but with statistically significant differences among phyla, habitats and temperature ranges that were mirrored by the number of KEGG level 3 functions in each genome. Particularly, archaea inhabiting extreme niches such as deep sea or hot springs characteristic with high local temperatures not only had significantly fewer total genes but also fewer KEGG level 3 functions. Otherwise, environments of higher complexity and diversity such as soils or sediments contain archaea with a larger functional potential that may allow them more options for intragenome competition or utilization of a wider range of nutrients.

### Functional redundancy

Most biogeochemical reactions are driven by a limited set of metabolic pathways that are found in a variety of microbial clades (Falkowski et al., 2008). In line with this observation, taxonomic diversity was found to correlate strongly with functional diversity (Rineau and Courty, 2011). Functions were classified into two groups as before (Starke et al., 2019, 2020a, 2020b): (i) highly redundant across species found in more than 90% of the species or (ii) unique to only a few found in less than 10% of the species. Here, intergenome redundancy was either high or low for roughly two thirds of all the KEGG functions; fewer than the 77.3% found in fungal genomes (Starke et al., 2019, 2020b). However, the presence of a higher share of functions of intermediate redundancy found in between 10% and 90% of the species suggested a less strict classification with more than two groups (Starke et al., 2019, 2020a, 2020b) that could be particularly important for organisms with smaller genomes such as archaea and bacteria. A set of functions present in a quarter of all archaea indicated that the presence of a driving phylotype or environment may drive intergenome redundancy. Indeed, the majority of functions (151/194) with an intergenome redundancy between 22% and 26% belonged to the phylum *Crenarchaeota*, the habitat hot springs and the temperature range of hyperthermophilic archaea, mainly affiliated with amino acid utilization, fermentation, methanogenesis and nucleic acid metabolism. The median intergenome redundancy was twice as high as found for fungi (Starke et al., 2019, 2020b), implying a higher share of functions shared among archaea on average. However, only half the gene copies (1.02 in archaea compared to 2.0 in fungi) were present, highlighting the close relationship between intergenome and intragenome redundancy. Indeed, the archaeal genomes revealed that low intergenome redundancy is generally related to high intragenome redundancy and vice versa. Presumably, every organism must choose between additional copies of functionally important genes or a higher number of different genes, especially in reduced genomes. Similarly to the pattern found in fungi (Starke et al., 2019, 2020b), functions belonging to the maintenance apparatus such as S-adenosylmethionine synthetase (EC 2.5.1.6, K00789) involved in the biosynthesis of amino acids were with both high intergenome and high intragenome redundancy allowing for more complex regulation of the gene, i.e. when more transcripts are needed. Otherwise, functions with low intergenome and low intragenome redundancy are highly specialized processes only performed by a few archaea such as the drug transporter MFS transporter, DHA1 family, multidrug resistance protein (K19578) found in the crenarchaeote *Thermofilum adornatus*.

### Archaeal contribution to the total microbiome functionality

The parametric approach estimated the archaeal contribution to the total microbiome functionality to roughly 4,200 KEGG level 3 functions; a magnitude less than predicted for both bacteria (Starke et al., 2020a) and fungi (Starke et al., 2019, 2020b, 2020a). The lower bound estimate of functional richness derived from the non-parametric approaches yielded roughly 3,000 KEGG level 3 functions. A plateau of functional richness with higher species richness made the predictions for archaea more reliable as the errors decrease with proximity to the asymptote (Thompson et al., 2003). In theory, more species must be sequenced until no new functions are found and the accumulation curve reaches the actual asymptote (Gotelli and Colwell, 2011) but, in practice, this approach is generally impossible due to the prohibitively large number of species that need to be sampled to reach an asymptote (Chao et al., 2009). In our meta-analysis, admittedly, the 417 genomes of distinct archaeal species only spanned three archaeal phyla with sequenced genomes from all 21 proposed archaeal phyla (Williams et al., 2017; Castelle and Banfield, 2018) and covered only a small part of the predicted taxonomic diversity in archaea; with databases containing up to 13,159 archaeal species (Yarza et al., 2014), the prediction of 5,000 archaeal genera (Amann and Rosselló-Móra, 2016) and the finding of 669 distinct archaeal species among 10,575 prokaryotic genomes (Zhu et al., 2019). Hence, the addition of genomes from novel archaeal species with potentially new KEGG level 3 functions could change both the parametric and the non-parametric estimates of functional richness. However, the differences in the estimates are likely not as tremendous as the potential differences in the estimates for both bacteria (Starke et al., 2020a) and fungi (Starke et al., 2019, 2020b, 2020a) as the accumulation curve already plateaued with 417 taxonomically distinct archaeal species. Noteworthy, it is unclear how well new functions are recovered in archaea. As there is notably less interest in archaea compared to bacteria, functional annotations might generally miss archaea-specific functions to a larger extent than bacteria-specific functions are missed in bacteria. As of today, our understanding of the contribution of archaea to the total microbiome functionality covers the majority of the KEGG level 3 functions but it could be that many as-yet unknown archaea-specific functions exist.

Our results suggest a limited contribution of archaea to the functional potential of the total microbiome, with a majority of archaeal functions already identified today. However, the existence of archaea-specific functions must be validated by novel and more sophisticated methods. The accumulation curve describing the increase of functional categories with the number of sequenced genomes in archaea was closer to the asymptote than in bacteria and fungi. This made the estimate of their contribution to the total microbiome functionality more precise, although it is still uncertain if the functional diversities of different domains can easily be compared. Noteworthy - and similar as in fungi - only a quarter of the genes in the archaeal genomes on average were affiliated with a KEGG level 3 function, clearly demonstrating the limitations of the annotation dataset as the prediction of functionality technically excluded three quarters of the entire functional potential. Different databases such as COG or Pfam could further improve our understanding of functional diversity, especially in archaea, as those covered three times more genes than KEGG. Still, different approaches and definitions of functionality could be necessary to estimate the actual number of functions.

## Materials and Methods

### Metadata collection of the total known archaeal microbiome functions

To explore the relationship between diversity and function and to compare genomes across archaeal phyla, habitats and temperature ranges, available genomes from archaea as taxonomic unit were downloaded from the integrated microbial genomes and microbiomes (IMG) of the Joint Genome Institute (JGI) on July 17^th^ 2020. One genome was randomly selected if a species had multiple sequenced genomes to yield taxonomically distinct archaeal species. For each genome, the gene counts for each function on the level 3 of KEGG Orthology (Kanehisa et al., 2016b, 2016a) as functional unit were retrieved. In total, our database comprised 417 completely annotated archaeal genomes with 2,835 KEGG functions (**Supplementary Table S1**). The sequencing status was denoted as “Draft” for one, “Permanent Draft” for 217 and “Finished” for 199 archaeal genomes. Only three genomes were available to describe psychrophilic archaea and were therefore excluded from further analysis. Intergenome redundancy was calculated as the number of KEGG functions covered by one randomly chosen species compared to the total number of functions in all species (Starke et al., 2020b). Intragenome redundancy or gene redundancy was estimated as average of genes per individual KEGG function in any one species (Starke et al., 2020b). The gene counts and KEGG functions per archaeal phylum, habitat and temperature range were retrieved as average with standard deviation from the database. To estimate the specific differences, both intergenome and intragenome redundancy were calculated for every phylum, habitat and temperature range as described for the total database above.

### Accumulation curves (AC)

Archaeal species were randomly added in intervals of one up to the maximum species richness of 417 with 1,000 random permutations per step using the function *specaccum* from the R package *vegan* (Oksanen et al., 2018). The AC of the database permutation was then fitted to a saturated (**Equation 1**) and an unsaturated model (**Equation 2**) with the critical point estimated by the term 3*A*_*f*_ (Čapek et al., 2016). Due to the plateauing shape of the AC, a logarithmic model was used as well. The fit of the models was compared by analysis of Akaike Information Criterion (AIC) (Bertrand et al., 2006) with a penalty per parameter set to *k* equals two. The total number of KEGG functions in archaea on Earth was predicted using a global species richness estimate of 13,159 archaeal species (Yarza et al., 2014) to calculate the potential maximum of KEGG functions via uncertainty propagation and Monte Carlo simulation of the function *predictNLS* in the R package *propagate* (Spiess, 2018). The non-parametric estimation of functional richness was calculated as Chao-1 (Chao, 1984, 1989). This method was developed to estimate the asymptotic species richness in a set of samples. Since our objective was to estimate the asymptotic functional richness, genomes took the role of samples and KEGG functions took the role of species in our analysis. Resampling and repeating computations for lower levels of sample accumulation generates a smooth curve of the estimations. A reliable estimator would reach its own asymptote before the species accumulation curve does (Gotelli and Colwell, 2001). To test whether this occurred in our dataset, Chao-1 was estimated using a random subset of every 50 picked archaeal genomes in the database starting with two species (**Equation 3**). Additionally, asymptotic functional richness was estimated using first order jackknife (jack-1) and the bootstrap “boot” methods with the function *specpool* in the R package *vegan* (Oksanen et al., 2018), to check if these two alternative methods yielded estimations comparable to the parametric and Chao-1 approaches.

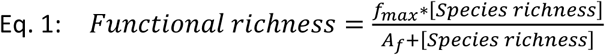

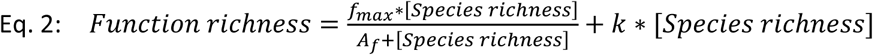

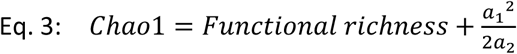

Here, *f*_*max*_ is the maximum functional richness, *A*_*f*_ the accretion rate of functions with an increasing number of species and *k* the constant of the additive term. Functions found only once or twice are indicated by a_1_ as singletons and a_2_ as doubletons, respectively.

## Acknowledgements

This work was supported by the Czech Science Foundation (20-02022Y). RS thanks Petr Capek for advice with the modelling.

## Author information

### Contributions

RS and PB designed the study. MLPF and RS performed the computational analysis. RS and IO modelled the data. The paper was written by MLPF, PB and RS, and reviewed by all authors. All authors approved the final version of the manuscript.

